# β-Adrenergic regulation of mGluR5-dependent Ca²⁺ signalling in reactive astrocytes

**DOI:** 10.64898/2026.11.25.769134

**Authors:** Matjaž Stenovec, Samo Pirnat, Julijan Vršnik, Klemen Dolinar, Alexei Verkhratsky, Robert Zorec

## Abstract

Astrocytes support neuronal function and brain homeostasis and undergo structural and functional remodelling under pathological conditions. Here, we compared astrocytes maintained in serum-containing medium, modelling a blood–brain barrier-compromised reactive state, with serum-free conditions that promote a more physiological arborised phenotype. We examined surface expression and signalling of metabotropic glutamate receptor 5 (mGluR5), β-adrenergic receptors (β-ARs), and pituitary adenylate cyclase-activating polypeptide type 1 receptor (PAC1) as well as expression of excitatory amino acid transporter 2 (EAAT2).

Live-cell immunolabelling combined with optophysiological measurements revealed that serum-exposed reactive astrocytes had substantially reduced surface mGluR5 expression and negligible mGluR5-dependent Ca^2+^ signalling, accompanied by increased expression of EAAT2. Adrenergic stimulation and sustained cAMP elevation restored surface mGluR5 expression and (S)-3,5-dihydroxyphenylglycine (DHPG)-evoked Ca^2+^ responses while reducing EAAT2 expression despite continued serum exposure. Reactive astrocytes also showed enhanced, prolonged β-AR-mediated cAMP responses that were largely uncoupled from changes in intracellular Ca^2+^.

These findings demonstrate coordinated remodelling of glutamatergic and β-adrenergic signalling in reactive astrocytes. The capacity to restore mGluR5-dependent signalling is retained and strongly regulated by cAMP, suggesting that neuromodulatory pathways can modulate astrocytic glutamate responsiveness independently of a complete reversal of the reactive phenotype.

## INTRODUCTION

Astrocytes maintain brain homeostasis throughout life, regulate extracellular environment, provide metabolic support to neurons ^1^, clear synaptic glutamate predominantly by the excitatory amino acid transporter 2 (EAAT2) ^2^ and detect activity of glutamatergic neurons by metabotropic glutamate receptors (in particular mGluR5) ^3^ that triggers intracellular Ca^2+^ signalling, which in turn modulates synaptic efficacy and plasticity ^4^. In response to acute injury or chronic disease, astrocytes mount reactive astrogliosis, leading to functional gain or loss ^5–9^. Understanding the signalling pathways that regulate EAAT2 and mGluR5 expression and localisation is essential for elucidating normal brain physiology and the pathogenesis of neurological disorders, as the local microenvironment strongly influences their expression and function. Because the exact composition of interstitial fluid surrounding individual cells *in vivo* is unknown and impossible to reproduce, an *in vitro* culture model provides a useful experimental approach. Astrocytes cultured in serum-containing medium adopt a reactive phenotype ^10^ as serum harbours a mixture of bioactive components that can trigger cellular activation, alter gene expression ^10,11^, and mimic several aspects of injury-induced astrocyte reactivity affecting the expression of mGluR5 and EAAT2. In contrast, astrocytes cultured in serum-free medium ^10,12^ retain a more realistic, *in vivo*–like phenotype^13^. This culture model minimises extrinsic activation signals, allowing intrinsic regulatory pathways to remain closer to their native state. Here, we used both culture models to investigate how an extrinsic factor such as serum affects baseline EAAT2 and mGluR5 expression and function in astrocytes. Notably, mGluRs have emerged as a promising therapeutic target for post-injury recovery because they are upregulated at lesion sites ^14^. However, it remains unclear how a reactive, serum-rich environment alters astrocytic mGluR5 and EAAT2 expression, and whether engaging intracellular signalling mechanisms can reverse these changes.

Beyond baseline differences, astrocytes respond dynamically to extracellular cues through cyclic adenosine monophosphate (cAMP)-dependent pathways. Activation of β-adrenergic receptors stimulates adenylyl cyclase ^15,16^, elevating intracellular cAMP ^17,18^ that activates protein kinase A (PKA), which phosphorylates cyclic AMP response element-binding protein (CREB) and initiates downstream transcriptional programs. PKA/CREB signalling enhances expression of the astrocytic glutamate transporter EAAT2 (GLT1) ^19^, and by extension could similarly affect mGluR5 expression. Thus, the β-AR–cAMP–PKA–CREB axis may represent a central mechanism by which astrocytes fine-tune both glutamate uptake capacity and receptor repertoire ^18,19^. By comparing serum-induced reactive (DMEM+) and serum-deprived arborised (NB+) astrocytes, we elucidated how culture milieu and differentiation state modulate astrocytic responsiveness to cAMP-elevating stimuli and the consequent transcriptional regulation of EAAT2 and mGluR5. Enhancing EAAT2 expression improves glutamate clearance and reduces excitotoxicity, whereas modulating mGluR5 fine-tunes Ca^2+^ signalling and synaptic support mechanisms.

Using Fluo-4 Ca^2+^ imaging combined with application of a selective mGluR5 agonist ((S)-3,5-dihydroxyphenylglycine; DHPG) and an antagonist (3-[2-(2-methyl-4-thiazolyl)ethynyl]-pyridine; MTEP), and live immunofluorescent labelling, we first assessed mGluR5 expression in serum-induced reactive astrocytes ^20^ and *in vivo*-like arborised astrocytes ^10,13^. Next, we tested whether prolonged increases in intracellular cAMP (by dbcAMP treatment or β-adrenergic receptor stimulation) restore reduced mGluR5 expression in serum-induced reactive astrocytes. We then examined EAAT2 surface density in both phenotypes by live immunolabeling and assessed how increased intracellular cAMP levels affect EAAT2 surface density in serum-induced reactive astrocytes. To increase translational relevance, we replicated key experiments in human astrocytes, confirming reduced mGluR5 surface expression in serum-induced reactive astrocytes and demonstrating restored mGluR5-dependent Ca^2+^ signalling after prolonged cAMP elevation. These findings suggest that cAMP-increasing treatments may restore astrocyte homeostatic phenotypes in pathological states with impaired blood-brain barrier function.

## MATERIALS AND METHODS

### Cell cultures

Primary astrocyte cultures were prepared from neocortices of 2–3-day-old Wistar rats (obtained from the Medical Experimental Centre at the Faculty of Medicine, University of Ljubljana, Slovenia) as previously described ^21,20^. Animal handling was in accordance with the European and Slovenian legislation (Official Gazette of the RS 38/13 and official consolidated text 21/18, 92/20, 159/21; UVHVVR, permit no. U34401-26/2025/). Cells were grown in DMEM+ medium (high-glucose [25 mM] Dulbecco’s modified Eagle’s medium (DMEM) supplemented with 10% fetal bovine serum, 1 mM sodium pyruvate, 2 mM L-glutamine, and 5 U/mL penicillin, 5 μg/mL streptomycin) in an atmosphere of 5% CO_2_/95% air. To purify isolated cells, sub-confluent cultures were shaken overnight (225 rpm), followed by medium replacement. Cells were then trypsinised, seeded onto poly-L-lysine-coated coverslips (50 μL suspension) and allowed to attach for 20 min at 37°C in 5% CO_2_/95% air. Cultures were maintained in DMEM+ or, after two washes with neurobasal medium (NB; Thermo Fisher Scientific, Waltham, MA, USA), in NB+ (NB supplemented with 2% B27, 5 ng/mL hbEGF, 2 mM GlutaMAX, 5 U/mL penicillin and 5 μg/mL streptomycin). Unless stated otherwise, chemicals were from Merck (Darmstadt, Germany) and were of the highest available purity.

A subset of experiments was conducted in foetal cortical human astrocytes (20 wk gestation, male, SC-1800, ScienCell, Carlsbad, CA, USA), which were obtained frozen, initially cultured for ∼1 week in Astrocyte Medium (AM, SC-1801, ScienCell), and subsequently in the same growth media as rat astrocytes (up to 3 weeks). We designate rat astrocytes cultured in DMEM+ or NB+ medium as DMEM+ or NB+ astrocytes, whereas human astrocytes cultured in the same media as hDMEM+ and hNB+, respectively.

### Live cell immunocytochemistry and fluorescence analysis

Surface-expressed EAAT2 and mGluR5 were immunolabelled in live astrocytes as described previously ^22^. Cells were first washed once with 3% BSA in PBS, incubated with 10% (v/v) goat serum in 3% (w/v) bovine serum albumin (BSA) in PBS for 1 h at 37°C to reduce the non-specific background staining, and then incubated either with rabbit polyclonal anti-EAAT2/GLT1 (NBP1-20136SS, 1:100; Novus Biologicals, Bio-Techne, Oxon, UK) or anti-mGluR5 (AGC-007, 1:100; Alomone, Jerusalem, Israel) diluted in 3% BSA in PBS for 45 min at 37°C. Subsequently, cells were washed three times with 3% BSA in PBS and incubated with the corresponding secondary antibody conjugated to Alexa Fluor 546 (1:600; Thermo Fisher Scientific) for 45 min at 37°C. Then, cells were washed three times with 3% BSA in PBS and once with extracellular solution, then transferred to the recording chamber on a confocal microscope (LSM 780, Carl Zeiss, Oberkochen, Germany), where they were observed with a plan-apochromatic oil-immersion objective (63×/NA 1.4). We obtained confocal z-stacked images using a 561-nm diode-pumped solid-state laser excitation, and band-pass filtered the fluorescence emission at 565–615 nm.

Surface EAAT2- and mGluR5-positive puncta were quantified per unit cell perimeter. Puncta were defined as clusters of ≥3 adjacent pixels (0.132 × 0.132 μm; minimum area, 0.052 µm^2^). Mean fluorescence intensity and apparent punctum area were quantified using ImageJ (NIH, MD, USA, http://rsbweb.nih.gov/ij/).

### Calcium imaging and analysis

Fluo-4 AM (5 μM; Thermo Fisher Scientific)-loaded cells were mounted on an LSM 780 confocal microscope and observed with a Plan-Apochromat 20×/NA 0.8 objective (Zeiss). We excited Fluo-4 with a 488-nm argon laser line and filtered the emission with a 495–565 nm band-pass filter. Time-lapse images were acquired every second for 5 min before and 10 min after bolus additions of L-Glu, 3-[2-(2-methyl-4-thiazolyl)ethynyl]-pyridine (MTEP), or (S)-3,5-dihydroxyphenylglycine (DHPG), to reach a final concentration of 1 mM, 1 µM and 10 μM, respectively. Baseline fluorescence was determined as the average fluorescence of the first 300 recordings (pre-stimulation).

### ICUE3 calibration, measurements and analysis

Astrocytes were transfected with plasmid encoding ICUE3 (0.8 µg/coverslip) ^23^ using FuGENE 6 (Promega Corporation, Madison, WI, USA) or Lipofectamine LTX (Thermo Fisher Scientific) and imaged 24–48 h later. Astrocyte-loaded coverslips were mounted in a chamber on an LSM 780 confocal microscope (Zeiss) and imaged using a 20×/0.8 NA air objective. ICUE3 was excited at 458 nm, and CFP and YFP emissions were recorded using a GaAsP detector with 464–499 nm and 516–534 nm band-pass filters, respectively.

To measure astrocytic [cAMP]_i_ we first calibrated ICUE3 (similarly as described in ^24^) and used the following equation: [cAMP] = EC_50_ × {(R – R_min_) / (R_max_ – R)}^1/n^, where EC_50_ is the cAMP concentration at which FRET response is half-maximal, n is Hill coefficient of the dose-response dependency, R_min_ is minimum CFP/YFP emission ratio (when adenylyl cyclase activity was inhibited by 10 min addition of 100 µM 2′,5′-dideoxyadenosine; DDA), and R_max_ is maximum CYP/YFP emission ratio when ICUE3 was saturated (after sequential 5 min additions of 8-Br-2′-*O*-Me-cAMP-AM reaching 2, 5, 20, 50 and 100 µM, respectively). After determining R_max_ and R_min_, we calculated [cAMP]_i_, either at rest or after cell stimulation.

L-Glu- and noradrenaline (NA)-evoked cAMP responses were monitored in DMEM+ and NB+ astrocytes by CFP/YFP time-lapse imaging (5-s intervals; 2 min baseline and 5 min after sequential addition of L-Glu (1 mM) and NA (100 µM)). Background-corrected fluorescence intensities were measured in individual-cell ROIs using ZEN 2010B (Zeiss), and cAMP responses were expressed as the CFP/YFP emission ratio after background subtraction (mean + 3×SD). An increase in [cAMP]_i_ was thus reflected by an increase in the CFP/YFP emission ratio.

To estimate the time constant (τ) of NA-evoked increases in [cAMP]_i_ in DMEM+ and NB+ astrocytes, we fitted a single-exponential increase to maximum function (f = y_0_+a × (1 − exp(−b × x)) to time-resolved changes in CFP/YFP emission ratio by custom-written MATLAB software taufit (MathWorks, Natick, MA, USA); τ = 1/b. The maximal amplitude (a_max_) of the [cAMP]_i_ increase was calculated as a quotient of the CFP/YFP emission ratio at the end of NA stimulation (averaged over the last 30 s) versus the resting CFP/YFP emission ratio before NA application (averaged over the last 30 s) and expressed as % increase or increase in CFP/YFP emission ratio.

### Fluorometric measurements of intracellular glutamate and glutamine

Intracellular glutamate and glutamine were quantified using the Glutamine/Glutamate-Glo™ Assay (Promega, Madison, WI, USA) according to the manufacturer’s protocol for cell lysates. Glutamate was measured directly, whereas total glutamate plus glutamine was determined after glutaminase treatment; glutamine was calculated by subtraction.

Primary rat cortical astrocytes were cultured in DMEM+ (10% FBS) or NB+ medium (B27 and 5 ng/mL hbEGF), seeded at 5 000 or 10 000 cells per well, and analysed at baseline (t_0_) or after 30-min exposure to 100 µM L-glutamate in extracellular solution containing 2 mM L-glutamine (t_30 min_), using separate plates. Cells were washed twice with PBS and lysed in 30 µL PBS plus 15 µL 0.3 N HCl; lysates were neutralised with 15 µL 450 mM Tris (pH 8.0), yielding 60 µL per well. For each lysate, duplicate 25-µL aliquots were incubated with glutaminase or enzyme-free buffer for 30–40 min at room temperature. Glutamate detection reagent (50 µL) was then added, followed by a 60-min incubation before luminescence measurement (Victor 3, PerkinElmer). Glutamate concentrations were determined from a calibration curve (assay range, approximately 0.05–50 µM), and glutamine concentrations by subtracting glutamate from total glutamate plus glutamine.

Metabolite levels were normalised to 10 000 cells based on independently determined proliferation, approximately two-fold in DMEM+ and 1.5-fold in NB+ culture. Accordingly, initial seeding densities of 5 000 and 10 000 cells corresponded to approximately 10 000 and 20 000 cells in DMEM+ cultures, and 7 500 and 15 000 cells in NB+ cultures, respectively.

### Statistical analysis

Data were analysed using SigmaPlot (Systat Software, San Jose, CA, USA) and are presented as mean ± SEM. Statistical significance was assessed using the Mann–Whitney U test or rank-based ANOVA followed by Dunn’s test, as appropriate. Intracellular L-Glu/L-Gln measurements (t_0_ vs. t_30 min_; separate plates) were compared using two-sided unpaired *t*-tests, with Welch’s correction when variances were unequal. *P* < 0.05 was considered significant.

## RESULTS

### L-glutamate selectively activates mGluR5 to evoke large Ca^2+^ increases in arborised, but not serum-induced reactive astrocytes from rat or human

First, we examined glutamate receptor-dependent Ca^2+^ signalling in serum-induced reactive (DMEM+) and in vivo-like arborised (NB+) astrocytes by measuring increases in [Ca^2+^]_i_ with non-ratiometric fluorescent Ca^2+^ indicator Fluo-4 ^12,13^; Fig. 1a, b). The application of L-Glu (1 mM) evoked minor, transient increases in Fluo-4 emission (Fig. 1a, middle and right; Fig. 1c, grey period) in a small fraction (28 out of 167; 17%) of DMEM+ astrocytes. In contrast, L-Glu evoked large, biphasic increases in [Ca^2+^]_i_ in all (77; 100%) NB+ astrocytes (Fig. 1b, middle and right; Fig. 1d, grey period) that also exhibited increased resting Ca^2+^ activity, as reported previously ^10,13^. To quantify increases in [Ca^2+^]_i_, we measured the peak Ca^2+^ amplitude (*p*) and the time-integrated increase in [Ca^2+^]_i_ (*S*, Δ*F*/*F*_0_×*s*) (Fig. 1c, d). At rest, little Ca^2+^ was mobilised in DMEM+ and NB+ astrocytes as revealed by small peak Ca^2+^ amplitudes (0.06 ± 0.00 versus 0.74 ± 0.11 Δ*F*/*F*_0_, *p*<0.001; Fig. 1e) and time-integrated Ca^2+^ activity (4.06 ± 0.34 versus 53.15 ± 6.62 Δ*F*/*F*_0_×*s*, *p*<0.001; Fig. 1f). Although the amount of free Ca^2+^ mobilised into the cytosol increased substantially upon L-Glu stimulation, it was substantially smaller in DMEM+ than NB+ astrocytes, as indicated by the peak amplitudes (1.15 ± 0.21 versus 3.10 ± 0.13 Δ*F*/*F*_0_, *p*<0.001, Fig. 1e) and by the time-integrated Ca^2+^ activity (36.92 ± 4.29 versus 416.31 ± 23.87 Δ*F*/*F*_0_×*s*, *p*<0.001, Fig. 1f).

**FIGURE 1.**
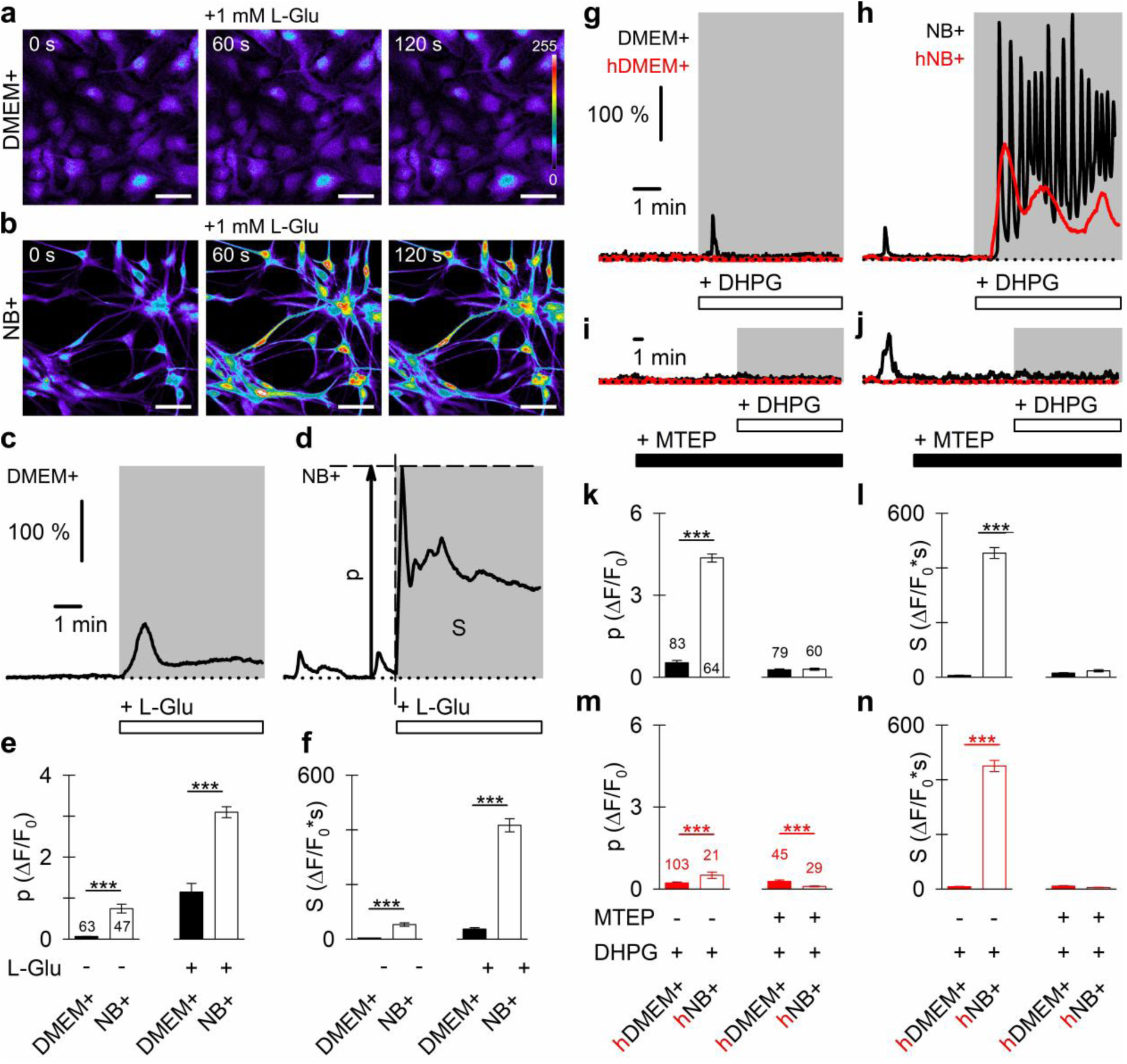
Arborised, but not serum-induced reactive astrocytes, respond to glutamate stimulation with large increases in [Ca^2+^]_i_. (**a, b**) Confocal micrographs of fluorescent Ca^2+^ indicator Fluo-4 loaded DMEM+ (**a**) and NB+ (**b**) astrocytes before (0 s, left) and after stimulation with 1 mM L-glutamate (L-Glu) to increase [Ca^2+^]_i_ (60 s (middle) and 120 s (right) after stimulus application). L-Glu evoked substantial increases in [Ca^2+^]_i_ in NB+, but not in DMEM+ astrocytes, as indicated by the pseudocoloured intensity scale (right, 0–255 intensity levels). Scale bars: 50 µm. (**c, d**) Time-resolved increases in [Ca^2+^]_i_ in DMEM+ (**c**) and NB+ (**d**) astrocytes were quantified by determining the peak [Ca^2+^]_i_ (*p*, mean±SEM) relative to the baseline fluorescence (grey horizontal line, *F*_0_) and the time-integrated [Ca^2+^]_i_ (*S*) at rest and after application of 1 mM L-Glu (white horizontal rectangle). The grey rectangular area indicates the L-Glu addition to astrocytes. (**e, f**) The peak (*p*; **e**) and the time-integrated calcium responses (*S*; **f**) in DMEM+ (black) and NB+ (white) at rest (**e, f, left**) and upon application of L-Glu (**e, f, right**). Note that L-Glu mobilised a substantial amount of Ca^2+^ into the cytosol of NB+, but not DMEM+ astrocytes. (**g, h**) Time-resolved increases in [Ca^2+^]_i_ in DMEM+ (**g**) and NB+ (**h**) rat (black) and human (red) astrocytes at rest and after application of 10 µM DHPG (100 µM in human astrocytes; white horizontal rectangle). Note that DHPG mobilised large amount of Ca^2+^ into the cytosol of NB+, but not DMEM+ rat astrocytes. (**i, j**) Time-resolved increases in [Ca^2+^]_i_ in DMEM+ (**i**) and NB+ (**j**) at rest, after application of 1 µM 3-[2-(2-methyl-4-thiazolyl)ethynyl]-pyridine (MTEP, black horizontal rectangle), and after application of 10 µM DHPG (white horizontal rectangle) to rat astrocytes. A selective antagonist of metabotropic glutamate receptor 5 completely blocks increases in [Ca^2+^]_i_ in both reactive and arborised astrocytes. (**k–n**) The peak (*p*; **k, m**) and the time-integrated calcium responses (*S*; **l, n**) in DMEM+ (black) and NB+ (white) at rest (**k, l, left**) and upon application of DHPG (**k, l, right**). The peak (*p*; **k, m**) and the time-integrated calcium responses (*S*) in DMEM+ (**l, n**; black or red) and NB+ (**l, n**; white or red) after application of MTEP (**i, j, left**), and subsequent application of DHPG (**i, j, right**). Note that DHPG did not mobilise Ca^2+^ into the cytosol of either NB+ or DMEM+ rat and human astrocytes pre-treated with MTEP. The grey rectangular area indicates the time of DHPG application (**g–j**). The numbers at the bottom of the bars indicate the number of cells analysed. \*\*\**P*<0.001 versus respective comparison (Mann-Whitney U test).

Distinct L-Glu-evoked increases in [Ca^2+^]_i_ in serum-induced reactive (DMEM+) and arborised (NB+) astrocytes suggested differential surface expression of glutamate receptors, presumably metabotropic glutamate receptor 5 (mGluR5) ^25^, which is essential in neurodevelopment ^26^, synaptic function and neuroprotection ^27^. Downregulation of mGluR5 may contribute to neurological and psychiatric disorders, while their excessive suppression impairs astrocyte function ^28^. To provide pharmacological evidences in support of differential mGluR5 expression, we applied selective mGluR5 agonist DHPG (10 µM) that evoked negligible increases in [Ca^2+^]_i_ in some (83 out of 180; 46%) DMEM+ astrocytes and substantial oscillatory increases in [Ca^2+^]_i_ in all (64 out of 64; 100%) NB+ astrocytes, as analytically confirmed by peak Ca^2+^ amplitudes (0.53 ± 0.07 versus 4.36 ± 0.14 Δ*F*/*F*_0_, *p*<0.001; Fig. 1k left) and the time-integrated Ca^2+^ activity (6.33 ± 1.11 versus 453.94 ± 19.94 Δ*F*/*F*_0_×*s*, *p*<0.001; Fig. 1l left). To provide further pharmacological evidence supporting mGluR5-dependent Ca^2+^ signalling, we pre-treated astrocytes with 1 µM MTEP (a negative allosteric modulator of mGluR5) and then stimulated them with 10 µM DHPG (Fig. 1i, j, k right, l right). Resting DMEM+ and NB+ astrocytes differed in basal Ca^2+^ activity (Fig. 1c–l) as observed earlier ^10,13^. After MTEP application, we detected no further increase in Ca^2+^ activity after the subsequent application of 10 µM DHPG. Our measurements confirmed DHPG-evoked Ca^2+^ activity by activation of mGluR5 (Fig. 1k right, l right) and indicated ample expression of mGluR5 in NB+, but not DMEM+ astrocytes. We further examined a clear and robust functional dichotomy in DHPG-evoked Ca^2+^ activity in human foetal cortical astrocytes cultured in the same media as rat astrocytes.

While primary cultures provide a powerful experimental platform for dissecting mechanisms of reactive astrogliosis, species-specific differences in astrocyte physiology necessitate validation in human cells, which more closely reflect the molecular and signalling landscape of the human brain and therefore offer greater translational relevance. To determine whether the remodelling of glutamatergic signalling identified in rat astrocytes extends across species, we reproduced the key optophysiological experiments in cultured human astrocytes (h-astrocytes) and examined mGluR5-dependent Ca^2+^ signalling in both cell types. Although 10 µM DHPG elicited only negligible increases in [Ca^2+^]ᵢ in h-astrocytes (not shown), one order of magnitude higher concentration (100 µM) failed to trigger Ca^2+^ activity in hDMEM+ astrocytes (Fig. 1g, red trace; Fig. 1m left, n left), but instead evoked larger, differential increases in [Ca^2+^]_i_ in the majority of hNB+ astrocytes (17 out of 21; 81%; Fig. 1h, red trace; Fig. 1n left). The differential response was analytically confirmed by larger peak Ca^2+^ amplitudes (0.22 ± 0.37 versus 0.50 ± 0.12 Δ*F*/*F*_0_, *p*<0.001; Fig. 1m left) and by increased time-integrated Ca^2+^ activity (8.43 ± 1.23 versus 449.74 ± 19.99 Δ*F*/*F*_0_×*s*, *p*<0.001; Fig. 1n left) in hNB+ astrocytes. Again, h-astrocytes pre-treated with MTEP (1 µM) failed to respond to DHPG (100 µM) (Fig. 1i, j), pharmacologically confirming mGluR5 involvement. MTEP alone did not elicit any Ca^2+^ activity (Fig. 1i, j, m right, n right).

### Adrenergic cAMP restores mGluR5-dependent Ca^2+^ signalling in reactive astrocytes

Large differences in DHPG-evoked Ca^2+^ activity in arborised (NB+) and serum-induced reactive (DMEM+) astrocytes (Fig. 1) suggested abundant mGluR5 expression in NB+ astrocytes. To provide independent structural evidence for differential mGluR5 expression, we immunofluorescently labelled mGluR5 at the surface of live rat astrocytes with primary anti-mGluR5 and the corresponding fluorescent secondary antibodies (Fig. 2a, b; similarly, as reported by ^22^). As expected, we found scarce mGluR5 immunolabelling in DMEM+, but ample in NB+ astrocytes (Fig. 2c, d). The quantification revealed ∼10-times less mGluR5-immunopositive puncta at surface of DMEM+ than NB+ astrocytes (0.07 ± 0.01 versus 0.65 ± 0.05 puncta/µm, respectively; Fig. 2e). The intensity and the apparent puncta area were also smaller in DMEM+ than in NB+ astrocytes (74.59 ± 1.18 versus 81.07 ± 0.53 a.u. (by 8%) and 0.10 ± 0.00 µm^2^ versus 0.26 ± 0.01 µm^2^ (by 59%), respectively; Fig. 2f, g). Prominent surface expression of mGluR5 in NB+ astrocytes corroborated fluorometric Ca²⁺ measurements (Fig. 1).

**FIGURE 2.**
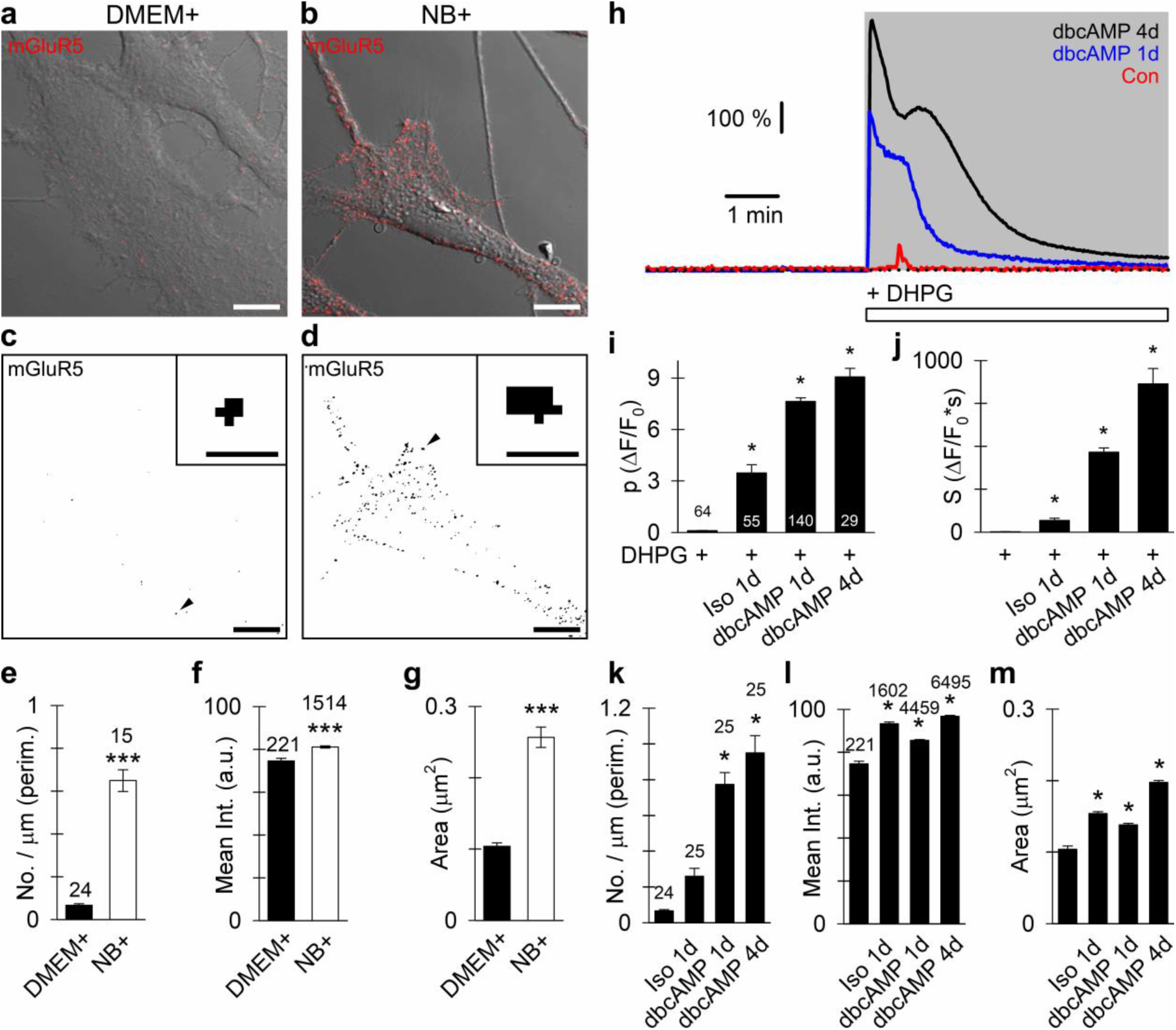
Metabotropic glutamate receptor 5 (mGluR5) is amply expressed at the surface of arborised but not serum-induced reactive astrocytes. (**a, b**) Superimposed differential interference contrast (DIC) images of serum (**a**; DMEM+) and arborised (**b**; NB+) astrocytes, and the corresponding confocal images of mGluR5-positive immunofluorescent puncta at the astrocyte surface (red). Note ample immunofluorescent puncta at the surface of NB+ (**b**), but not DMEM+ astrocytes (**a**). (**c, d**) Binarised mask display (black) of mGluR5-positive puncta in DMEM+ (**c**) and NB+ (**d**) astrocytes, respectively. Insets (**c, d**) display a magnified view of an individual mGluR5-positive puncta (indicated by an arrow) in both astrocytes. Scale bars: 10 μm (**large images, c, d**) and 1 μm (**insets, c, d**). (**e–g**) Number of mGluR5-positive puncta per micrometre of cell perimeter (**e**), mean intensity (**f**; in arbitrary units, a.u.), and surface area (**g**; in µm^2^, mean±SEM) of immunofluorescent mGluR5-positive puncta at the surface of DMEM+ (black) and NB+ (white) astrocytes. The numbers above the bars indicate the number of cells (**e**) or immunofluorescent mGluR5-positive puncta analysed (**f, g**), respectively. \*\*\**P* < 0.001 (Mann-Whitney U test). (**h**) Pharmacological treatments that increase [cAMP]_i_ restore DHPG-evoked calcium signalling in serum-induced reactive astrocytes. Time-resolved increase in [Ca^2+^]_i_ at rest and after application of 10 µM DHPG (white horizontal rectangle) in non-treated control DMEM+ astrocyte (red) and astrocyte treated for 1 day (1d) with 1 mM dbcAMP (blue) or for 4 days (4d; black). The grey rectangular area indicates the DHPG addition time. (**i, j**) The peak (*p*; **i**) and the time-integrated calcium responses (*S*; **j**) evoked by application of DHPG to non-treated DMEM+ astrocytes and DMEM+ astrocytes treated with dbcAMP for 1 day (dbcAMP, 1d) or 4 days (dbcAMP, 4d) or treated with Iso for 1 day (Iso, 1d; black). DHPG mobilised a large amount of Ca^2+^ in DMEM+ astrocytes treated with agents that increase [cAMP]_i_, but not in non-treated DMEM+ controls. The numbers at the bottom of the bars indicate the number of cells analysed. \**P*<0.05 versus non-treated DMEM+ controls (ANOVA on ranks followed by Dunn’s test). (**k–m**) Number of mGluR5-positive puncta per µm of cell perimeter (**k**), mean intensity (**l**; in arbitrary units, a.u.), and surface area (**m**; in µm^2^, mean±SEM) of mGluR5-positive puncta at the surface of non-treated DMEM+ astrocytes and astrocytes treated with Iso for 1 day, dbcAMP for 1 day, and dbcAMP for 4 days, respectively. The numbers above the bars indicate the number of cells (**e, k**) or immunofluorescent mGluR5-positive puncta analysed (**f, g, l, m**), respectively. \**P* < 0.05, \*\*\**P* < 0.001 (Mann-Whitney U test).

As demonstrated previously ^13^, arborised astrocytes exhibit increased resting intracellular cAMP concentration ([cAMP]ᵢ), a key modulator of astrocytic morphological complexity ^17,29–31^. As cAMP signalling confers an *in vivo*-like phenotype to astrocytes ^32^, we next examined whether treatments that increase [cAMP]_i_ in serum-induced reactive (DMEM+) astrocytes reconstitute mGluR5 surface expression and restore DHPG-evoked Ca^2+^ activity. Serum-induced reactive astrocytes were grown in serum-containing medium and treated for 1 day with either 100 µM isoprenaline (Iso), a β-adrenergic agonist, or 1 mM dbcAMP for 1 or 4 days, then examined for DHPG-evoked Ca^2+^ activity. At rest, negligible Ca^2+^ was mobilised into the cytosol of DMEM+ astrocytes treated either with dbcAMP (Fig. 2h, red trace) or Iso (not shown), similarly to non-treated DMEM+ controls (Fig. 1). Application of DHPG evoked substantial increases in [Ca^2+^]_i_ in Iso- and dbcAMP-treated DMEM+ astrocytes, but not in non-treated controls (Fig. 2h). The peak Ca^2+^ amplitudes (Δ*F*/*F*_0_) of 0.53 ± 0.07 (DMEM+ controls), 3.47 ± 0.47 (DMEM+ & Iso, 1 day; *p*<0.05), 7.63 ± 0.21 (DMEM+ & dbcAMP, 1 day; *p*<0.05) and 9.06 ± 0.50 (DMEM+ & dbcAMP, 4 days; *p*<0.05; Fig. 2i), and the time-integrated Ca^2+^ activity (Δ*F*/*F*_0_×*s*) of 6.33 ± 1.11 (DMEM+), 67.64 ± 12.81 (DMEM+ & Iso, 1 day; *p*<0.05; Fig. 2j), 464.84 ± 25.07 (DMEM+ & dbcAMP, 1 day; *p*<0.05) and 863.42 ± 89.01 (DMEM+ & dbcAMP, 4 days; *p*<0.05) revealed restored mGluR5-dependent Ca^2+^ activity. Prolonged 4-day dbcAMP treatment appeared the most effective, while modest 1-day Iso treatment was the least effective. Despite these differences, all treatments restored DHPG-evoked Ca^2+^ activity above levels observed in non-treated DMEM+ controls (Fig. 2h–j). Notably, the temporal profiles of DHPG-evoked Ca^2+^ activity differed. In Iso- and dbcAMP-treated DMEM+ astrocytes, the initial peak in [Ca^2+^]_i_ was followed by a rapid or delayed progressive decrease in [Ca^2+^]_i_ (peak–plateau-type response, Fig. 2h) in the majority (55-91%) of examined astrocytes, whereas in NB+ astrocytes oscillatory increases in [Ca^2+^]_i_ (Fig. 1d) predominated (94%).

As treatments that increase [cAMP]_i_ restore DHPG-evoked Ca^2+^ activity in serum-induced reactive astrocytes, we also immunocytochemically examined mGluR5 expression in treated astrocyte groups. As expected, all treatments restored mGluR5 surface expression. The detailed quantification revealed ∼3-4-times more mGluR5-positive puncta per unit length (µm^−1^) in DMEM+ astrocytes treated with dbcAMP than with Iso, however all treatments that increased [cAMP]_i_ gave rise to an increase (*p*<0.05; ANOVA on ranks followed by Dunn’s test) in mGluR5-positive puncta when compared to non-treated astrocytes, (0.07 ± 0.01, DMEM+; 0.26 ± 0.04 puncta/µm; DMEM+ & Iso, 1 day, 0.77 ± 0.07, DMEM+ & dbcAMP, 1 day and 0.95 ± 0.09, DMEM+ & dbcAMP, 4 days; Fig. 2k). The puncta intensity was smaller in non-treated controls when compared to astrocytes treated with Iso or dbcAMP; 74.59 ± 1.18, 93.22 ± 0.84, 85.51 ± 0.40, 96.74 ± 0.39 a.u. (Fig. 2l). The apparent puncta area was also smaller in non-treated astrocytes when compared to astrocytes treated with Iso or dbcAMP; 0.10 ± 0.00, and 0.15 ± 0.00, 0.14 ± 0.00, 0.19 ± 0.00 µm^2^, respectively (Fig. 2m). The differences in the intensity and mGluR5-positive puncta area may indicate differences in the copy number of mGluR5 inside the receptor cluster (hot spots) at the cell surface. Live immunocytochemical labelling confirmed reconstituted mGluR5 surface expression in serum-induced reactive astrocytes following treatments that increase [cAMP]_i_. Surface expression of excitatory amino acid transporter 2 (EAAT2) is augmented in serum-induced reactive astrocytes and downregulated by prolonged increase in intracellular cAMP concentration

Glutamate uptake and its metabolic conversion to glutamine are considered canonical homeostatic functions and are expected to be impaired in serum-induced reactive astrocytes ^33–35^. Thus, we immunocytochemically examined surface expression of excitatory amino acid transporter 2 (EAAT2) in both astrocytes (Fig. 3a, b). EAAT2-positive immunofluorescent puncta appeared more abundant in DMEM+ than NB+ astrocytes (Fig. 3c, d), which was analytically confirmed by quantifying puncta number per unit length (µm^−1^) of cell perimeter (0.62 ± 0.08 versus 0.29 ± 0.05 puncta/µm, respectively; Fig. 3e). Both, the puncta intensity and the apparent area were larger in DMEM+ than NB+ astrocytes (99 ± 1 versus 96 ± 1 a.u. (by 3%) and 0.33 ± 0.00 µm^2^ versus 0.28 ± 0.01 µm^2^ (by 16%), respectively; Fig. 3f, g).

**FIGURE 3.**
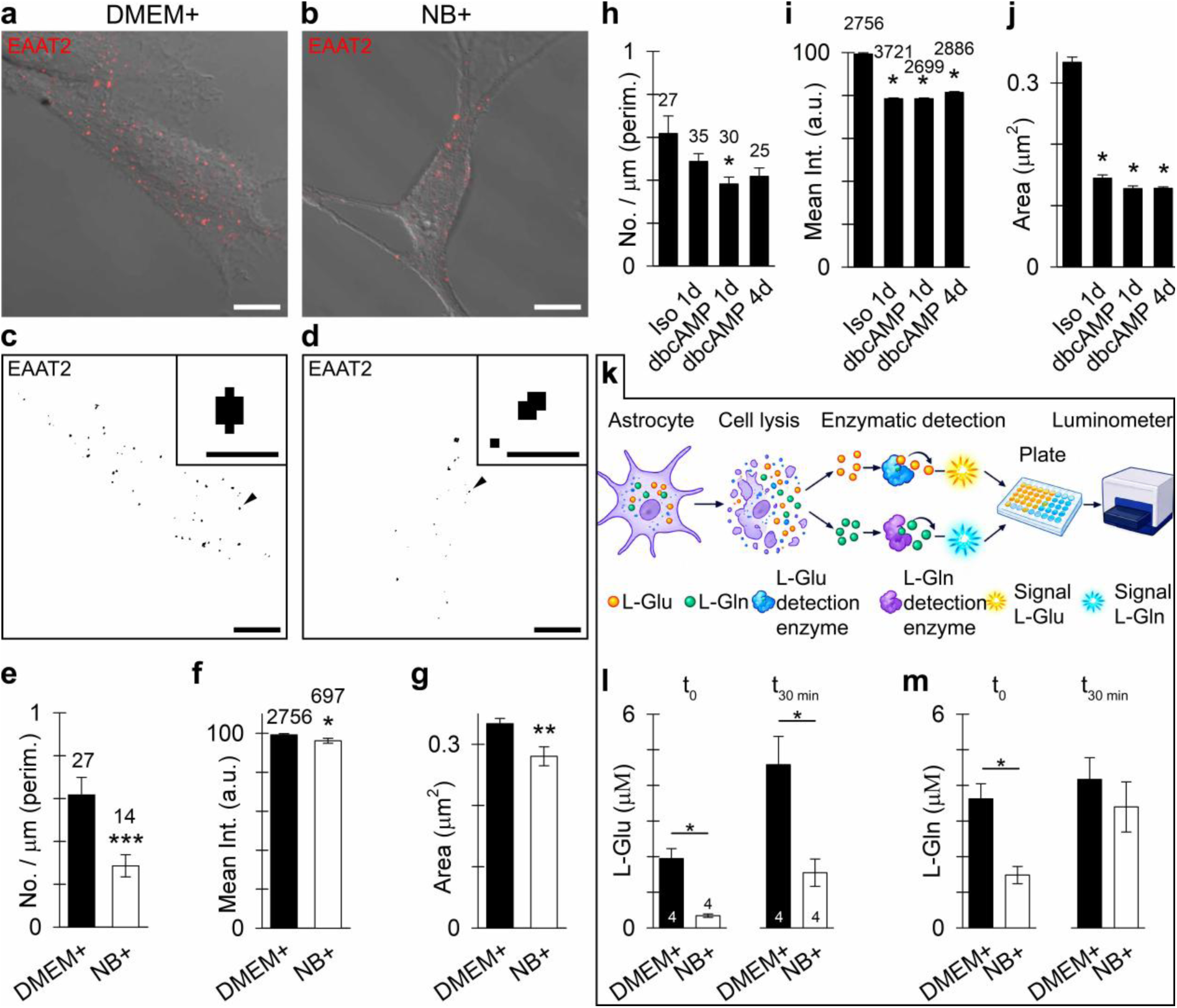
Reduced surface expression of excitatory amino acid transporter 2 (EAAT2) in arborised astrocytes. (**a, b**) Superimposed, differential interference contrast (DIC) images of serum (**a**; DMEM+) and arborised (**b**; NB+) astrocytes, and the corresponding confocal images of EAAT2-positive immunofluorescent puncta at the astrocyte surface (red). Note the more numerous EAAT2-positive puncta at the surface of DMEM+ (**a**) than NB+ (**b**) astrocytes. (**c, d**) Binarised mask display (black) of EAAT2-positive puncta in DMEM+ (**c**) and NB+ (**d**) astrocytes, respectively. Insets (**c, d**) display a magnified view of an individual EAAT2-positive puncta (indicated by an arrow) in both astrocytes. Scale bars: 10 μm (**large images, c, d**) and 1 μm (**insets, c, d**). (**e–g**) Number of EAAT2-positive puncta per µm of cell perimeter (**e**), mean intensity (**f**; in arbitrary units, a.u.), and surface area (**g**; in µm^2^, mean±SEM) of immunofluorescent EAAT2-positive puncta at the surface of DMEM+ (black) and NB+ (white) astrocytes. The numbers above the bars indicate the number of cells (**e**) or immunofluorescent EAAT2-positive puncta analysed (**f, g**), respectively. \**P* < 0.05, \*\**P* < 0.01, \*\*\**P* < 0.001 (Mann-Whitney U test). (**h–j**) Serum-induced reactive astrocytes express less EAAT2 upon treatments that increase [cAMP]_i_. Number of EAAT2-positive puncta per micrometre of cell perimeter (**h**), mean intensity (**i**; in arbitrary units, a.u.), and the apparent puncta area (**j**; in µm^2^, mean±SEM) in non-treated and DMEM+ astrocytes treated with 100 µM Iso for 1 day or with 1 mM dbcAMP for 1 or 4 days (1d or 4d). The numbers above the bars indicate the number of cells (**h**) or immunofluorescent EAAT2-positive puncta analysed (**i, j**), respectively. \**P* < 0.05 (ANOVA on ranks followed by Dunn’s test). (**k**) Basic principle of the Glutamine/Glutamate-Glo™ assay used to quantify intracellular L-Glu and L-Gln in cell lysates through enzyme-coupled luminescence detection. (**l, m**) Intracellular L-Glu (**l**) and L-Gln (**m**) concentrations (mean±SEM; n = 4 normalised wells per condition) in DMEM+ (black) and NB+ astrocytes (white) measured at t_0_ and t_30 min_, 30 min after the cell exposure to exogenous 100 µM L-Glu. Only statistically significant within-group comparisons at baseline (t_0_) and after 30 min (t_30 min_) are displayed (\**P* < 0.05; two-sided unpaired t-test).

Surprisingly, serum-induced reactive astrocytes displayed higher EAAT2 surface expression than arborised astrocytes, even though the arborised phenotype more closely resembles mature, homeostatic *in vivo*-like astrocytes ^34,36^. Next, we asked whether treatments that increase [cAMP]_i_ alter the expression of EAAT2 in DMEM+ astrocytes as dbcAMP treatment causes astrocytes to differentiate, by stimulating specific gene expression ^37^. Treatments with Iso for 1 day or dbcAMP for 1 or 4 days downregulated EAAT2 surface expression. The EAAT2-positive puncta/µm of cell perimeter decreased modestly from 0.62 ± 0.08 (DMEM+) to 0.49 ± 0.04 (DMEM+ & Iso, 1 day), 0.38 ± 0.03 (DMEM+ & dbcAMP, 1 day), and 0.42± 0.04 (DMEM+ & dbcAMP, 4 days; *P*<0.05; Fig. 3h, ANOVA on ranks followed by Dunn’s test), whereas the intensity and the apparent puncta area decreased substantially (*P*<0.05, ANOVA on ranks followed by Dunn’s test) in all treated groups. The puncta intensity decreased from 99 ± 1 (DMEM+) to 79 ± 0 a.u. (DMEM+ & Iso, 1 day), 79 ± 0 (DMEM+ & dbcAMP, 1 day; *P*<0.05) and 81 ± 0 (DMEM+ & dbcAMP, 4 days; *P*<0.05; Fig. 3i), while the apparent EAAT2 puncta area decreased from 0.33 ± 0.00 to 0.14 ± 0.00 (*P*<0.05), 0.13 ± 0.00 (*P*<0.05) and 0.13± 0.00 µm^2^ (*P*<0.05; Fig. 3j), respectively.

To provide functional evidence for differential surface density of EAAT2 and glutamate handling in serum-induced reactive and arborised astrocytes, we conducted populational fluorometric measurements of L-Glu//L-Gln using the Glutamine/Glutamate-Glo Assay to quantify intracellular L-Glu and L-Gln in both types of astrocytes at baseline (t_0_) and following a 30-min challenge with 100 µM L-Glu (t_30 min_; Fig. 3k) under continuous presence of extracellular L-Gln (2 mM). In DMEM+ astrocytes, intracellular L-Glu increased substantially from 1.94 ± 0.29 to 4.58 ± 0.79 µM (*P* = 0.020), whereas L-Gln increased minutely from 3.62 ± 0.43 to 4.17 ± 0.61 µM (*P* = 0.493). In NB+ astrocytes, L-Glu increased from 0.34 ± 0.05 to 1.55 ± 0.38 µM (*P* = 0.020), and L-Gln from 1.48 ± 0.24 to 3.40 ± 0.70 µM (*P* = 0.042; two-sided unpaired t-tests). We normalised values for both metabolites to 10 000 cells. Over 30 min, the corresponding apparent net intracellular accumulation rates were 5.27 and 1.09 pmol/min/10 000 cells for L-Glu and L-Gln, respectively, in DMEM+ astrocytes, and 2.41 and 3.83 pmol/min/10 000 cells in NB+ astrocytes. Thus, the higher surface abundance of EAAT2 in DMEM+ astrocytes translated to a larger absolute intracellular L-Glu pool at rest (t_0_) and net L-Glu accumulation over 30 min (t_30 min_; Fig. 3l, m), whereas NB+ astrocytes displayed a relatively greater increase in the L-Gln pool upon glutamate challenge (Fig. 3l, m).

### β-adrenergic receptor-dependent cAMP production is larger, but delayed in serum-induced reactive astrocytes

Because prolonged Iso treatment of serum-induced reactive astrocytes altered mGluR5 and EAAT2 expression (Figs. 1–3), we next examined whether β-adrenergic receptor–mediated cAMP signalling also differs between the two astrocyte types. Notably, arborised astrocytes exhibit increased resting intracellular cAMP ([cAMP]ᵢ) ^13^, consistent with the established role of increased [cAMP]ᵢ in promoting astrocyte arborisation ^17,29–31^. As serum-free astrocytes cultured in astrocyte-defined medium expressed detectable mRNA for mGluR3 ^25^ that potentially translates to protein, which may mediate L-Glu–dependent cAMP responses and (paradoxically) increase cAMP in astrocytes in the presence of extracellular Ca^2+^ ^38^, we examined L-Glu–stimulated cAMP production in serum-induced reactive and arborised astrocytes. Astrocytes were transfected with the plasmid encoding FRET-based cAMP nanosensor ICUE3 (Fig. 4a, b), related to ICUE2 that reports [cAMP]_i_ with EC_50_ ∼12.5 µM ^39^. As EC_50_ was not determined in ICUE3, we first provided cAMP calibration data enabling conversion of CFP/YFP emission ratio values into [cAMP]_i_. For calibration, we first treated ICUE3-expressing DMEM+ astrocytes (47 cells from 2 cultures) for 10 min with 100 µM DDA, then sequentially for 5 min with 2, 5, 20, 50, and 100 µM 8-Br-2′-O-Me-cAMP-AM (cAMP for simplicity). During the course of treatment, the mean CFP/YFP emission ratio changed from 0.44 (resting; Suppl. Fig. 1a) to 0.43 (100 µM DDA), 0.50 (2 µM cAMP), 0.58 (5 µM cAMP), 0.73 (20 µM cAMP), 0.73 (50 µM cAMP) and 0.76 (100 µM cAMP; Suppl. Fig. 1b). Data points (mean±SEM) were fitted by the four-parameter logistic curve of the form: f = min + (max-min)/(1 + (x/EC_50_)^(-Hillslope)2^), with min = 0.42 ± 0.03, max=0.78 ± 0.05, EC_50_ = 6.02 ± 2.79, and Hill slope = 1.00 ± 0.40 (Suppl. Fig. 1c).

**FIGURE 4.**
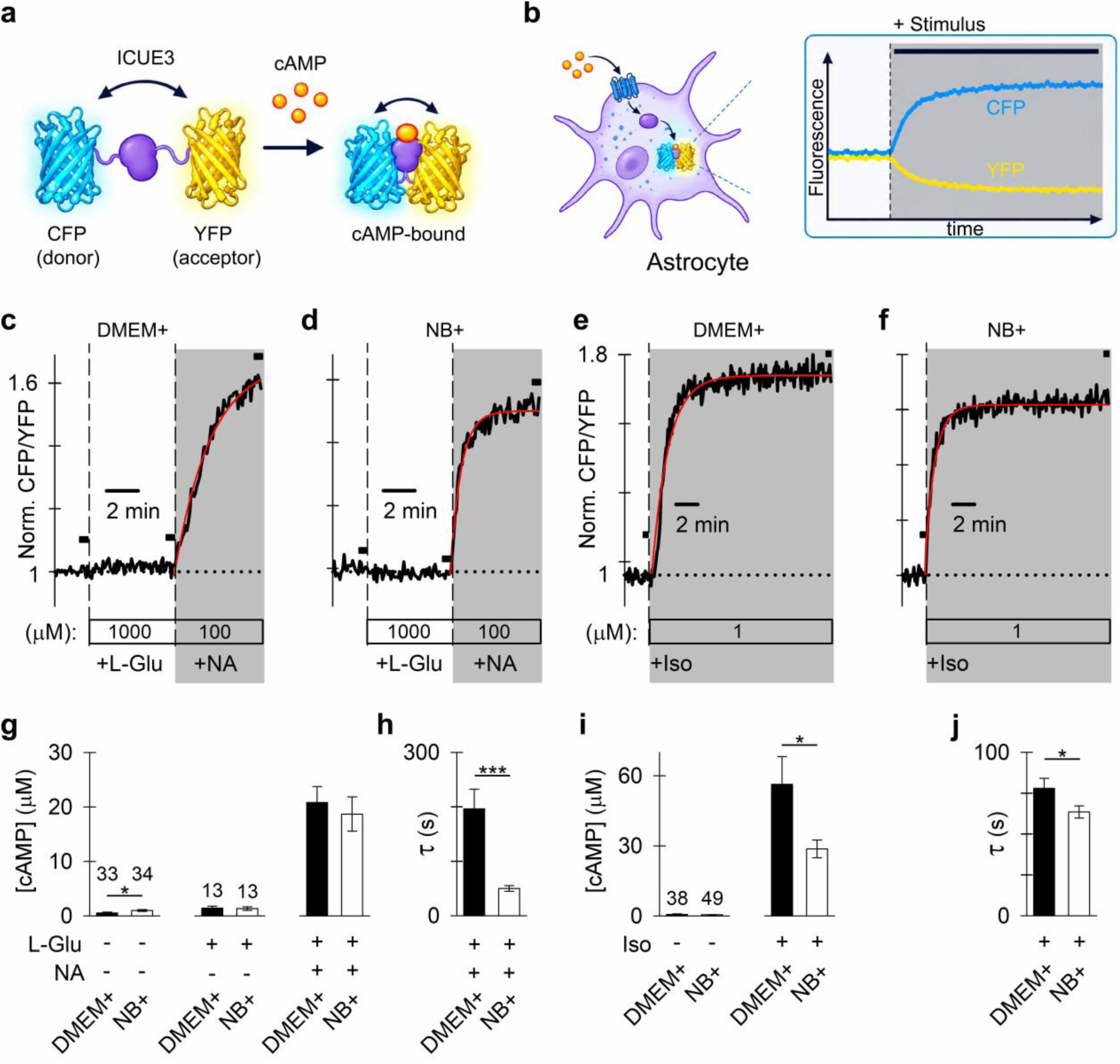
Noradrenaline and isoprenaline evoke faster increases in [cAMP]_i_ in arborised astrocytes. (**a, b**) Schematic overview of the ICUE3 biosensor design (**a**) and experimental workflow to quantify receptor-mediated changes in intracellular cAMP levels in living astrocytes by fluorescence ratio imaging (**b**). (**c–f**) Time-dependent changes in normalised CFP/YFP emission ratio measured in DMEM+ (**c, e**) and NB+ (**d, f**) astrocytes that were sequentially stimulated with 1 mM L-Glu and then with 100 µM NA (**c, d**) or with 1 µM Iso (**e, f**). The monophasic increase in CFP/YFP emission ratio evoked by NA indicates an increase in [cAMP]_i_ that was fitted by a single-exponential increase to maximum function (red; f = y_0_ + a × (1 − exp(−b × x)), where τ = 1/b; see Materials and Methods). (**g, i**) Mean (±SEM) [cAMP]_i_ (in µM) at rest (**g, i**), after stimulation with 1 mM L-Glu (**g**) and after stimulation with 100 µM NA (**g**) or with 1 µM Iso (**i**); all measured within 30 s (small black rectangles above traces on **c– f**) at the end of the stimulation period. L-Glu did not increase [cAMP]i, whereas NA evoked a large increase in DMEM+ and NB+ astrocytes with similar maximal amplitude (**g**) but different kinetics (**h**), and 1 µM Iso evoked larger increases in [cAMP]_i_ in DMEM+ astrocytes (**i**). (**h, j**) The time constant (τ) of the normalised CFP/YFP emission ratio increase in astrocytes stimulated by NA (**h**) and Iso (**j**). Note faster increase in [cAMP]_i_ in NB+ than DMEM+ astrocytes. \**P* < 0.05, \*\*\**P* < 0.001 (Mann-Whitney U test).

After calibrating ICUE3, we examined L-Glu and NA-evoked increases in [cAMP]_i_ in both astrocytes (Fig. 4c, d). As previously described ^13^the resting [cAMP]_i_ (pooled data) was lower in DMEM+ than in NB+ astrocytes (0.55 ± 0.14 µM versus 0.99 ± 0.17 µM; Fig. 4g left). Application of 1 mM L-Glu did not change the normalised CFP/YFP ratio (Fig. 4c, d, g middle), whereas stimulation with 100 µM NA evoked a large, time-dependent increase in [cAMP]_i_ in both astrocytes (Fig. 4c, d, g right). The maximum amplitude of NA-evoked increase in [cAMP]_i_ was similar (Fig. 4g right), whereas the time constant (τ) of increase in [cAMP]_i_ was greater in DMEM+ than in NB+ astrocytes (196.4 ± 0.3 s versus 50.1 ± 5.2 s, respectively; Fig. 4h).

To verify that different kinetics of NA-evoked increase in [cAMP]_i_ is not due to an interference of concurrently active Ca^2+^ and cAMP signalling pathways ^40–42^, we next examined cAMP increases evoked by 1 µM isoprenaline, a specific β-adrenergic receptor agonist, with no concurrent mobilisation of Ca^2+^ in astrocytes ^43^. The application of 1 µM Iso evoked large time-dependent increases in [cAMP]_i_ in DMEM+ and NB+ astrocytes (Fig. 4e, f) with ∼ twice larger maximum amplitude (56.4 ± 11.9 µM versus 28.7 ± 3.8 µM; Fig. 4i right) and larger time constant (τ; slower kinetics) in DMEM+ than in NB+ astrocytes (79.8 ± 7.7 s versus 55.9 ± 5.1 s, respectively; Fig. 4j). It appears that preceding (L-Glu-evoked) or concurrent (NA-evoked) mobilisation of Ca^2+^ in astrocyte cytosol does not account for faster cAMP production in arborised astrocytes. These results likely indicate a phenotype-specific autonomous difference in the cAMP production pathway.

To further examine differences in cAMP production in both astrocytes, we applied full-length pituitary adenylate cyclase-activating polypeptide (PACAP1-38) to DMEM+ and NB+ astrocytes (Fig. 5a–d). PACAP1-38 (10 nM) robustly elevated intracellular cAMP in both DMEM+ and NB+ astrocytes either in transient (Fig. 5a, b) or sustained manner (Fig. 5c, d), producing peak [cAMP]_i_ of 9.1 ± 2.5 µM versus 11.1 ± 3.0 µM (Fig. 5e right), and the time constant (τ) of increase in [cAMP]_i_ (rise kinetics), 98.4 ± 20.0 s versus 91.8 ± 22.2 s, in DMEM+ and NB+ astrocytes, respectively (Fig. 5f). Although neither parameter differed significantly between culture conditions, PACAP1-38 elicited a striking redistribution of response type. In DMEM+ astrocytes, 69 % of cells exhibited a transient increase in [cAMP]_i_ and only 31 % generated a sustained plateau, whereas in NB+ astrocytes only 21 % of responses were transient and 79 % sustained (χ² = 10.19, df = 1, *P*=0.001). Although

**FIGURE 5.**
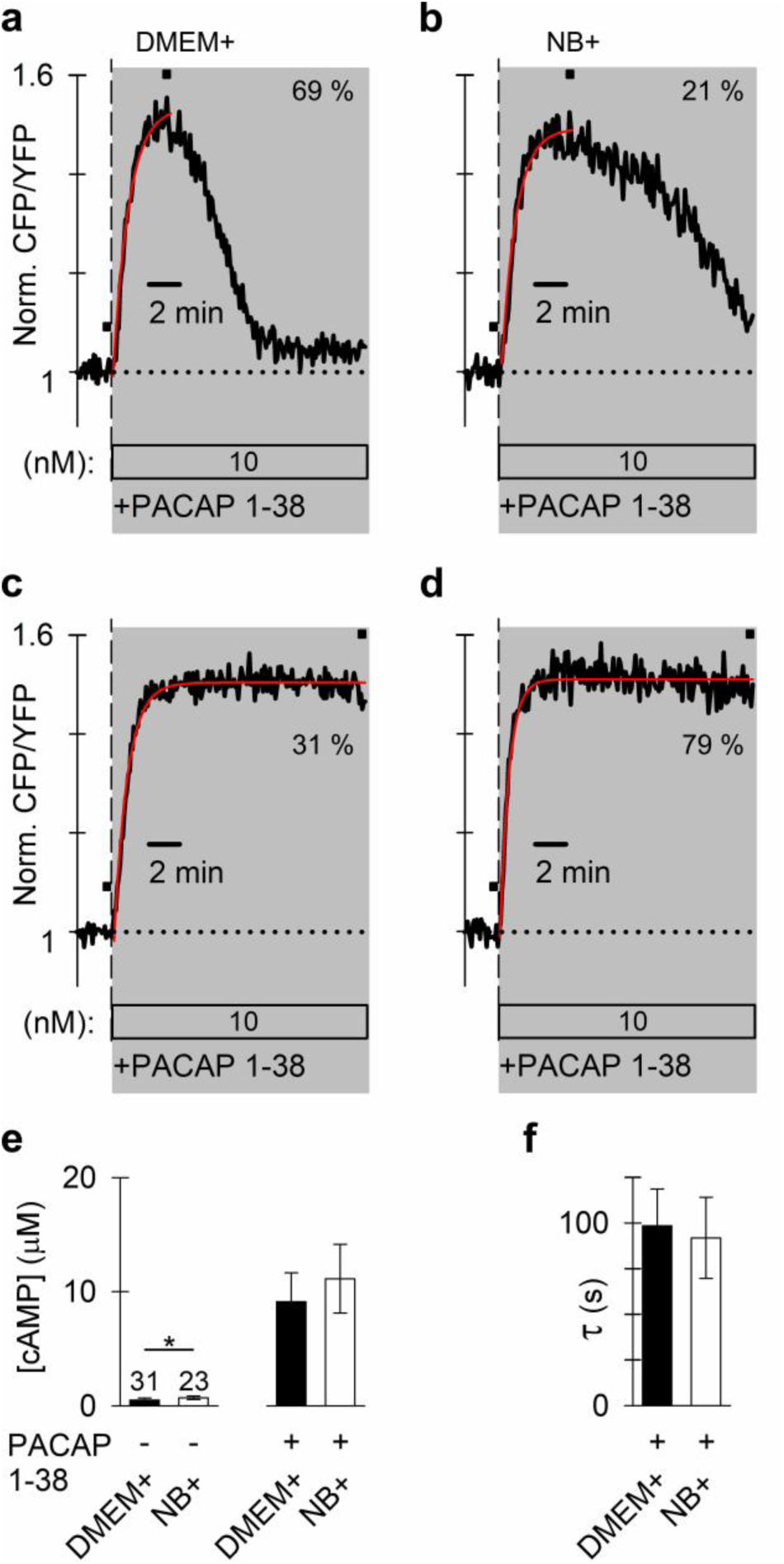
Full-length pituitary adenylate cyclase-activating polypeptide (PACAP1-38)-evoked increases in [cAMP]_i_ are similar in amplitude and kinetics, but different in relative proportion of transient and sustained cAMP responses in arborised and serum-induced reactive astrocytes. (**a–d**) Time-dependent changes in normalised CFP/YFP emission ratio evoked by PACAP1-38 (10 nM) in DMEM+ (**a, c**) and NB+ (**b, d**) astrocytes. Note that the relative proportion of PACAP1-38-evoked transient (**a, b**) and sustained (**c, d**) increases in normalised CFP/YFP emission differed in DMEM+ and NB+ astrocytes. Increases in normalised CFP/YFP emission ratio were fitted (red line) as described in the legend to Fig. 4 c–f (see also Materials and Methods). (**e**) Mean (±SEM) [cAMP]_i_ (in µM) at rest (**e left**) and after stimulation with 10 nM PACAP1-38 (**e right**) measured in 30 s (small black rectangles above traces on **a–d**) at the end or the peak increase in the given period. In both astrocytes, PACAP1-38 evoked increases in [cAMP]_i_ with similar maximal amplitude (**e right**) and kinetics (**f**). \**P* < 0.05 (Mann-Whitney U test).

PACAP1-38 stimulation preserved absolute cAMP amplitude and kinetics, prolonged cAMP signalling in arborised NB+ astrocytes suggested a context-dependent bias in second-messenger encoding that may underlie divergent downstream gene-regulatory and functional outcomes. Thus, dissimilar proportions in PACAP1-38-evoked responses indicate that both receptor engagement and astrocyte state shape distinct second-messenger dynamics.

## DISCUSSION

We investigated glutamate-dependent Ca^2+^ and cAMP signalling in astrocytes cultured in serum-containing medium (DMEM+), promoting a reactive phenotype associated with blood–brain barrier disruption ^44^, or serum-free neurobasal medium (NB+), promoting an *in vivo*-like arborised phenotype ^10,12,13^. We examined how these conditions remodel mGluR5 and EAAT2 function, and whether elevated intracellular cAMP can modify their expression and function.

### Culture-dependent regulation and cAMP-mediated rescue of mGluR5

Our Ca^2+^ imaging and live-cell immunocytochemistry revealed abundant presence of surface mGluR5 in arborised astrocytes but markedly reduced expression in serum-exposed reactive astrocytes (Figs. 1, 2). This finding is consistent with previous studies showing that astrocytic mGluR5 expression depends strongly on the extracellular environment. Serum-free and growth factor-supplemented conditions promote mGluR5 expression, whereas serum-containing conditions can preserve its mRNA without detectable protein production ^25,45–47^. In our study, supplementation of serum-free neurobasal medium with heparin-binding EGF-like growth factor (hbEGF; 5 ng/mL) supported astrocyte survival, process extension and arborisation, together with robust mGluR5-dependent signalling. Conversely, hbEGF alone did not reverse the flat morphology associated with serum-containing culture conditions ^48,10,12,13^. The mechanism by which hbEGF promotes mGluR5 expression remains unresolved but may involve transcriptional regulation, mRNA stabilisation or both.

Inflammatory and injury-associated signals, including interleukin-1β, thrombin and factors released by reactive microglia, can suppress astrocytic mGluR5 expression ^47,49,50^. Because mGluR5 activation can also suppress astrocytic glutamate transporter expression, its downregulation may represent an adaptive response to inflammatory stress, although its consequences are likely context-dependent ^45^. Of note, serum exposure attenuated mGluR5-dependent Ca^2+^ signalling in human foetal and newborn rat astrocytes, suggesting this response is conserved across species. These findings raise the possibility that serum-associated or injury-related environmental cues shift astrocytes away from an mGluR5-enriched signalling state relevant to neuroglial communication and development.

Prolonged elevation of intracellular cAMP restored surface mGluR5 expression and DHPG-evoked Ca^2+^ responses in serum-exposed astrocytes (Fig. 2), consistent with a role for sustained cAMP signalling in promoting astrocytic differentiation ^32^. This effect may involve cAMP/PKA-dependent transcriptional regulation, receptor trafficking or both ^51,52^, although our experiments do not distinguish between these mechanisms. Nevertheless, the recovery of mGluR5-dependent signalling demonstrates that serum-associated suppression is not necessarily irreversible and that astrocyte signalling properties remain amenable to pharmacological modulation. This finding provides a basis for investigating alternative routes of cAMP activation, including PACAP receptor signalling, as potential means of modulating astrocyte function when conventional β-adrenergic signalling is compromised.

### Phenotype-dependent regulation of EAAT2 and glutamate handling

Despite their less reactive morphology, lower GFAP expression and elevated basal intracellular Ca^2+^ and cAMP levels, arborised NB+ astrocytes exhibited lower EAAT2 surface density than serum-exposed DMEM+ astrocytes ^13^. *In situ* (rodent brain slices), EAAT2 is predominantly localised in clusters along peri-synaptic processes that enwrap active glutamatergic synapses ^53–55^, while its expression positively correlates with synaptic activity ^56^ and is upregulated by increased glutamate release from neurons ^54,57^. Given its major contribution to glutamate uptake in the brain, the higher EAAT2 surface abundance in DMEM+ astrocytes may represent an adaptive response to serum-associated stress, potentially limiting excitotoxic exposure. However, surface expression alone does not establish transporter activity or demonstrate neuroprotection.

In NB+ astrocytes, several mechanisms may contribute to the lower EAAT2 surface density. The vitamin A-containing B27 supplement used in serum-free medium may influence retinoid signalling, as retinoic acid can repress EAAT2 expression and promote its protein kinase C-dependent internalisation ^58^. In addition, reduced GFAP expression may affect transporter trafficking ^59,60^ as astrocytes from GFAP-deficient mice exhibit impaired EAAT2 delivery to the plasma membrane following PKA activation ^61^. These mechanisms provide plausible explanations for the phenotype-dependent differences but remain to be tested in our culture model. Conversely, the preservation of EAAT2 expression under serum-associated conditions is consistent with evidence that thrombin selectively downregulates EAAT1 while sparing EAAT2 in primary rat cortical astrocytes ^36^.

The supplementary L-glutamate (L-Glu) and L-glutamine (L-Gln) measurements provide functional context for these differences. Following exposure to 100 µM glutamate, DMEM+ astrocytes exhibited a larger net increase in intracellular L-Glu than NB+ astrocytes over 30 min (158 versus 72 pmol/10 000 cells), corresponding to apparent net accumulation rates of 5.27 and 2.41 pmol/min/10 000 cells, respectively. This pattern is consistent with greater net glutamate accumulation in serum-exposed astrocytes and agrees with their higher EAAT2 surface density. Nevertheless, intracellular L-Glu accumulation reflects the balance between uptake, metabolism, release and other processes; it does not directly quantify EAAT2-specific transport flux.

Conversely, the net increase in intracellular L-Gln was greater in NB+ than in DMEM+ astrocytes (115 versus 33 pmol/10 000 cells). The ratio of apparent L-Gln to L-Glu net accumulation rates was approximately 1.59 in NB+ astrocytes and 0.21 in DMEM+ astrocytes. Because 2 mM L-Gln was continuously present extracellularly at both measurement time points, the difference in intracellular L-Gln accumulation cannot be attributed to a change in extracellular glutamine availability between t_0_ and t_30 min_. However, intracellular L-Gln reflects the combined effects of synthesis, uptake, efflux and metabolic consumption, and therefore does not directly measure conversion of glutamate into glutamine. Together, these measurements indicate distinct patterns of intracellular glutamate and glutamine accumulation. DMEM+ astrocytes accumulate more glutamate, whereas NB+ astrocytes show a greater relative increase in the glutamine pool.

Prolonged dbcAMP treatment (1–4 days) reduced EAAT2 surface expression in serum-exposed astrocytes (Fig. 3), indicating that elevated cAMP does not uniformly regulate astrocytic membrane proteins. This result contrasts with reports that prolonged dbcAMP treatment increases GLT1 expression in primary rat astrocytes, including following mechanical injury ^37^. Such differences may reflect variation in culture conditions, treatment duration and the distinction between transporter gene expression and surface localisation. Thus, cAMP-dependent regulation of EAAT2 appears context-dependent, potentially involving changes in expression and trafficking that need not occur in parallel.

### Astrocyte phenotype shapes β-adrenergic and PACAP-dependent cAMP signalling

Despite contrasting mGluR5 and EAAT2 expression, β-adrenergic receptor- and PACAP1–38-dependent cAMP signalling remained functional in both astrocyte phenotypes, although response kinetics differed (Figs. 4, 5). Iso induced a steeper cAMP rise in NB+ than in DMEM+ astrocytes, whereas PACAP1–38 elicited more sustained responses in NB+ cells. These differences may reflect phenotype-dependent regulation of receptor coupling, adenylyl cyclase activity or phosphodiesterase-mediated cAMP degradation, potentially modulated by intracellular Ca^2+^ levels. Although the underlying mechanisms remain unresolved, distinct cAMP kinetics may differentially engage downstream effectors, including PKA and exchange protein directly activated by cAMP (EPAC), and thereby influence astrocyte function and plasticity ^62–64^. However, our data do not establish a causal link between cAMP dynamics and mGluR5 or EAAT2 surface expression.

Preserved PACAP1–38 responsiveness also suggests an alternative route for cAMP activation ^64^ when β-adrenergic signalling is compromised. Whether PACAP can compensate functionally for impaired β-adrenergic input or restore receptor and transporter function remains to be investigated.

Overall, our findings reveal coordinated but non-uniform remodelling of glutamate handling and intracellular signalling across astrocyte phenotypes. Serum-associated reactivity combines reduced mGluR5-dependent Ca^2+^ signalling with increased EAAT2 surface expression, potentially coupling diminished neuromodulatory responsiveness to enhanced glutamate clearance. Conversely, NB+ astrocytes exhibit prominent mGluR5-dependent responses and distinct glutamate and glutamine accumulation profiles. These findings underscore the importance of defining astrocyte states by functional properties rather than morphology or reactivity markers alone. The reversibility of mGluR5 suppression following prolonged cAMP elevation further identifies cAMP signalling as a potential means of modifying astrocyte phenotype. Future studies should establish the molecular basis and functional consequences of receptor-specific cAMP dynamics, particularly in pathological models involving blood–brain barrier disruption and neurological disease. Such investigations may identify strategies that preserve the protective functions of reactive astrocytes while restoring deficient neuroglial communication.

## ACKNOWLEDGEMENTS

We acknowledge financial support from the Slovenian Research and Innovation Agency (ARIS), Core Research Program P3-310 Cell Physiology, ARIS grants J4-60077, J3-50104, J1-70029, N3-0470, I0-0034 [Celica], I0-0048 [CIPKeBiP] and I0-0022 [MRIC UL]. We thank Ms E. Rus for expert technical assistance in preparing growth media, experimental solutions, and maintaining astrocyte cultures.

## DATA AVAILABILITY STATEMENT

The data are available upon reasonable request from the corresponding author.

Supplementary information accompanies the manuscript on the Experimental & Molecular Medicine website (http://www.nature.com/emm/).

## Author contributions

M.S. generated the idea, conceived the study, designed and performed the confocal imaging experiments, performed image analysis and quantification, constructed figures and wrote the manuscript. S.P. and J.V. multiplied plasmids, transfected and labelled cells, performed image analysis and quantification, and wrote the manuscript. S.P. and K.D. performed population fluorometric measurements of L-Glu/L-Gln and analysed the data; R.Z. contributed to experimental design, wrote and revised the manuscript, and provided funding (P3 310). A.V. conceived the study, secured funding, and supervised the project. All authors discussed the results, critically revised the manuscript and approved the final version.

## REFERENCES

1 Verkhratsky, A. & Nedergaard, M. Physiology of Astroglia. Physiol Rev 98, 239–389 (2018). 10.1152/physrev.00042.2016

2 Rothstein, J. D. et al. Knockout of glutamate transporters reveals a major role for astroglial transport in excitotoxicity and clearance of glutamate. Neuron 16, 675–686 (1996). https://doi.org/S0896-6273(00)80086-0 [pii] 10.1016/s0896-6273(00)80086-0

3 Pasti, L., Volterra, A., Pozzan, T. & Carmignoto, G. Intracellular calcium oscillations in astrocytes: a highly plastic, bidirectional form of communication between neurons and astrocytes in situ. J Neurosci 17, 7817–7830 (1997). 10.1523/JNEUROSCI.17-20-07817.1997

4 Henneberger, C. & Rusakov, D. A. Synaptic plasticity and Ca2+ signalling in astrocytes. Neuron Glia Biol 6, 141–146 (2010). https://doi.org/S1740925X10000153 [pii] 10.1017/S1740925X10000153

5 Pekny, M. & Pekna, M. Astrocyte reactivity and reactive astrogliosis: costs and benefits. Physiol Rev 94, 1077–1098 (2014). https://doi.org/94/4/1077 [pii] 10.1152/physrev.00041.2013

6 Sofroniew, M. V. Molecular dissection of reactive astrogliosis and glial scar formation. Trends Neurosci 32, 638–647 (2009). 10.1016/j.tins.2009.08.002

7 Verkhratsky, A. et al. Astrocytes in human central nervous system diseases: a frontier for new therapies. Signal Transduct Target Ther 8, 396 (2023). 10.1038/s41392-023-01628-9

8 Verkhratsky, A. et al. Curing the brain: in search for new astrocyte-specific therapies. Exp Mol Med 58, 1086–1127 (2026). 10.1038/s12276-026-01712-4

9 Verkhratsky, A., Zorec, R. & Parpura, V. Stratification of astrocytes in healthy and diseased brain. Brain Pathol 27, 629–644 (2017). 10.1111/bpa.12537

10 Wolfes, A. C. et al. A novel method for culturing stellate astrocytes reveals spatially distinct Ca2+ signaling and vesicle recycling in astrocytic processes. J Gen Physiol 149, 149–170 (2017). 10.1085/jgp.201611607

11 Zhang, Y. et al. An RNA-sequencing transcriptome and splicing database of glia, neurons, and vascular cells of the cerebral cortex. J Neurosci 34, 11929–11947 (2014). 34/36/11929 [pii] 1860-14 [pii] 10.1523/JNEUROSCI.1860-14.2014

12 Wolfes, A. C. & Dean, C. Culturing In Vivo-like Murine Astrocytes Using the Fast, Simple, and Inexpensive AWESAM Protocol. J Vis Exp (2018). 10.3791/56092

13 Pirnat, S. et al. Astrocyte arborization enhances Ca(2+) but not cAMP signaling plasticity. Glia 69, 2899–2916 (2021). https://doi.org/GLIA24076 [pii] 10.1002/glia.24076

14 Spampinato, S. F., Copani, A., Nicoletti, F., Sortino, M. A. & Caraci, F. Metabotropic Glutamate Receptors in Glial Cells: A New Potential Target for Neuroprotection? Front Mol Neurosci 11, 414 (2018). 10.3389/fnmol.2018.00414

15 Horvat, A., Zorec, R. & Vardjan, N. Adrenergic stimulation of single rat astrocytes results in distinct temporal changes in intracellular Ca(2+) and cAMP-dependent PKA responses. Cell Calcium 59, 156–163 (2016). 10.1016/j.ceca.2016.01.002

16 Shao, Y. & Sutin, J. Expression of adrenergic receptors in individual astrocytes and motor neurons isolated from the adult rat brain. Glia 6, 108–117 (1992). 10.1002/glia.440060205

17 Vardjan, N., Kreft, M. & Zorec, R. Dynamics of β-adrenergic/cAMP signaling and morphological changes in cultured astrocytes. Glia 62, 566–579 (2014). 10.1002/glia.22626

18 Zhou, Z., Ikegaya, Y. & Koyama, R. The Astrocytic cAMP Pathway in Health and Disease. Int J Mol Sci 20 (2019). ijms20030779 [pii] ijms-20-00779 [pii] 10.3390/ijms20030779

19 Karki, P. et al. cAMP response element-binding protein (CREB) and nuclear factor kappaB mediate the tamoxifen-induced up-regulation of glutamate transporter 1 (GLT-1) in rat astrocytes. J Biol Chem 288, 28975–28986 (2013). https://doi.org/S0021-9258(20)48871-4 [pii] M113.483826 [pii]10.1074/jbc.M113.483826

20 McCarthy, K. D. & de Vellis, J. Preparation of separate astroglial and oligodendroglial cell cultures from rat cerebral tissue. J Cell Biol 85, 890–902 (1980).

21 Schwartz, J. P. & Wilson, D. J. Preparation and characterization of type 1 astrocytes cultured from adult rat cortex, cerebellum, and striatum. Glia 5, 75–80 (1992). 10.1002/glia.440050111

22 Bozic, M., Verkhratsky, A., Zorec, R. & Stenovec, M. Exocytosis of large-diameter lysosomes mediates interferon gamma-induced relocation of MHC class II molecules toward the surface of astrocytes. Cell Mol Life Sci 77, 3245–3264 (2020). https://doi.org/10.1007/s00018-019-03350-8 10.1007/s00018-019-03350-8 [pii]

23 DiPilato, L. M. & Zhang, J. The role of membrane microdomains in shaping beta2-adrenergic receptor-mediated cAMP dynamics. Mol Biosyst 5, 832–837 (2009). 10.1039/b823243a

24 Borner, S. et al. FRET measurements of intracellular cAMP concentrations and cAMP analog permeability in intact cells. Nat Protoc 6, 427–438 (2011). https://doi.org/nprot.2010.198 [pii]10.1038/nprot.2010.198

25 Janssens, N. & Lesage, A. S. Glutamate receptor subunit expression in primary neuronal and secondary glial cultures. J Neurochem 77, 1457–1474 (2001). 10.1046/j.1471-4159.2001.00369.x

26 Danjo, Y. et al. Transient astrocytic mGluR5 expression drives synaptic plasticity and subsequent chronic pain in mice. J Exp Med 219 (2022). https://doi.org/213089 [pii]jem.20210989 [pii]10.1084/jem.20210989

27 Vanzulli, I. & Butt, A. M. mGluR5 protect astrocytes from ischemic damage in postnatal CNS white matter. Cell Calcium 58, 423–430 (2015). https://doi.org/S0143-4160(15)00113-X [pii]10.1016/j.ceca.2015.06.010

28 de Lima, I. B. Q. et al. Blockade of mGluR5 in astrocytes derived from human iPSCs modulates astrocytic function and increases phagocytosis. Front Immunol 14, 1283331 (2023). 10.3389/fimmu.2023.1283331

29 Koyama, Y., Ishibashi, T., Hayata, K. & Baba, A. Endothelins modulate dibutyryl cAMP-induced stellation of cultured astrocytes. Brain Res 600, 81–88 (1993). 10.1016/0006-8993(93)90404-b

30 Schiweck, J., Eickholt, B. J. & Murk, K. Important Shapeshifter: Mechanisms Allowing Astrocytes to Respond to the Changing Nervous System During Development, Injury and Disease. Front Cell Neurosci 12, 261 (2018). 10.3389/fncel.2018.00261

31 Won, C. L. & Oh, Y. S. cAMP-induced stellation in primary astrocyte cultures with regional heterogeneity. Brain Res 887, 250–258 (2000).

32 Paco, S., Hummel, M., Pla, V., Sumoy, L. & Aguado, F. Cyclic AMP signaling restricts activation and promotes maturation and antioxidant defenses in astrocytes. BMC genomics 17, 304 (2016). 10.1186/s12864-016-2623-4

33 Campbell, S. C. et al. Potassium and glutamate transport is impaired in scar-forming tumor-associated astrocytes. Neurochem Int 133, 104628 (2020). https://doi.org/S0197-0186(19)30363-8 [pii]10.1016/j.neuint.2019.104628

34 Escartin, C. et al. Reactive astrocyte nomenclature, definitions, and future directions. Nat Neurosci 24, 312–325 (2021). 10.1038/s41593-020-00783-4

35 Jiang, R., Diaz-Castro, B., Looger, L. L. & Khakh, B. S. Dysfunctional Calcium and Glutamate Signaling in Striatal Astrocytes from Huntington’s Disease Model Mice. J Neurosci 36, 3453–3470 (2016). 36/12/3453 [pii]3693-15 [pii]10.1523/JNEUROSCI.3693-15.2016

36 Piao, C. et al. Thrombin decreases expression of the glutamate transporter GLAST and inhibits glutamate uptake in primary cortical astrocytes via the Rho kinase pathway. Exp Neurol 273, 288–300 (2015). https://doi.org/S0014-4886(15)30091-1 [pii]10.1016/j.expneurol.2015.09.009

37 Eng, D. L., Lee, Y. L. & Lal, P. G. Expression of glutamate uptake transporters after dibutyryl cyclic AMP differentiation and traumatic injury in cultured astrocytes. Brain Res 778, 215–221 (1997). https://doi.org/S0006-8993(97)01093-7 [pii]10.1016/s0006-8993(97)01093-7

38 Moldrich, R. X., Aprico, K., Diwakarla, S., O’Shea, R. D. & Beart, P. M. Astrocyte mGlu(2/3)-mediated cAMP potentiation is calcium sensitive: studies in murine neuronal and astrocyte cultures. Neuropharmacology 43, 189–203 (2002). https://doi.org/S0028390802001119 [pii]10.1016/s0028-3908(02)00111-9

39 Violin, J. D. et al. beta2-adrenergic receptor signaling and desensitization elucidated by quantitative modeling of real time cAMP dynamics. J Biol Chem 283, 2949–2961 (2008). https://doi.org/S0021-9258(20)55543-9 [pii]10.1074/jbc.M707009200

40 Ahuja, M., Jha, A., Maleth, J., Park, S. & Muallem, S. cAMP and Ca(2)(+) signaling in secretory epithelia: crosstalk and synergism. Cell Calcium 55, 385–393 (2014). https://doi.org/S0143-4160(14)00022-0 [pii]10.1016/j.ceca.2014.01.006

41 Antoni, F. A. Interactions between intracellular free Ca2+ and cyclic AMP in neuroendocrine cells. Cell Calcium 51, 260–266 (2012). https://doi.org/S0143-4160(11)00242-9 [pii]10.1016/j.ceca.2011.12.013

42 Bruce, J. I., Straub, S. V. & Yule, D. I. Crosstalk between cAMP and Ca2+ signaling in non-excitable cells. Cell Calcium 34, 431–444 (2003). https://doi.org/S0143416003001507 [pii]10.1016/s0143-4160(03)00150-7

43 Horvat, A. et al. Ca(2+) as the prime trigger of aerobic glycolysis in astrocytes. Cell Calcium 95, 102368 (2021). https://doi.org/S0143-4160(21)00022-1 [pii]10.1016/j.ceca.2021.102368

44 Zamanian, J. L. et al. Genomic analysis of reactive astrogliosis. J Neurosci 32, 6391– 6410 (2012). 10.1523/JNEUROSCI.6221-11.2012

45 Aronica, E. et al. Expression and functional role of mGluR3 and mGluR5 in human astrocytes and glioma cells: opposite regulation of glutamate transporter proteins. Eur J Neurosci 17, 2106–2118 (2003). https://doi.org/2657 [pii]10.1046/j.1460-9568.2003.02657.x

46 Miller, S., Bridges, R. J. & Cotman, C. W. Stimulation of phosphoinositide hydrolysis by trans-(+/-)-ACPD is greatly enhanced when astrocytes are cultured in a serum-free defined medium. Brain Res 618, 175–178 (1993). https://doi.org/0006-8993(93)90443-Q [pii]10.1016/0006-8993(93)90443-q

47 Miller, S., Sehati, N., Romano, C. & Cotman, C. W. Exposure of astrocytes to thrombin reduces levels of the metabotropic glutamate receptor mGluR5. J Neurochem 67, 1435–1447 (1996). 10.1046/j.1471-4159.1996.67041435.x

48 Tsugane, M., Nagai, Y., Kimura, Y., Oka, J. & Kimura, H. Differentiated astrocytes acquire sensitivity to hydrogen sulfide that is diminished by the transformation into reactive astrocytes. Antioxid Redox Signal 9, 257–269 (2007). 10.1089/ars.2007.9.257

49 Aronica, E., Gorter, J. A., Rozemuller, A. J., Yankaya, B. & Troost, D. Interleukin-1 beta down-regulates the expression of metabotropic glutamate receptor 5 in cultured human astrocytes. J Neuroimmunol 160, 188–194 (2005). https://doi.org/S0165-5728(04)00446-1 [pii]10.1016/j.jneuroim.2004.11.015

50 Tilleux, S., Berger, J. & Hermans, E. Induction of astrogliosis by activated microglia is associated with a down-regulation of metabotropic glutamate receptor 5. J Neuroimmunol 189, 23–30 (2007). https://doi.org/S0165-5728(07)00213-5 [pii]10.1016/j.jneuroim.2007.06.011

51 Francesconi, A. & Duvoisin, R. M. Role of the second and third intracellular loops of metabotropic glutamate receptors in mediating dual signal transduction activation. J Biol Chem 273, 5615–5624 (1998). https://doi.org/S0021-9258(18)67818-4 [pii]10.1074/jbc.273.10.5615

52 Lee, J. H. et al. Calmodulin dynamically regulates the trafficking of the metabotropic glutamate receptor mGluR5. Proc Natl Acad Sci U S A 105, 12575–12580 (2008). https://doi.org/0712033105 [pii]4343 [pii]10.1073/pnas.0712033105

53 Minelli, A., Barbaresi, P., Reimer, R. J., Edwards, R. H. & Conti, F. The glial glutamate transporter GLT-1 is localized both in the vicinity of and at distance from axon terminals in the rat cerebral cortex. Neuroscience 108, 51–59 (2001). https://doi.org/S0306-4522(01)00375-X [pii]10.1016/s0306-4522(01)00375-x

54 Poitry-Yamate, C. L., Vutskits, L. & Rauen, T. Neuronal-induced and glutamate-dependent activation of glial glutamate transporter function. J Neurochem 82, 987– 997 (2002). https://doi.org/1075 [pii]10.1046/j.1471-4159.2002.01075.x

55 Zhou, J. & Sutherland, M. L. Glutamate transporter cluster formation in astrocytic processes regulates glutamate uptake activity. J Neurosci 24, 6301–6306 (2004). 24/28/6301 [pii]10.1523/JNEUROSCI.1404-04.2004

56 Swanson, R. A. et al. Neuronal regulation of glutamate transporter subtype expression in astrocytes. J Neurosci 17, 932–940 (1997). 10.1523/JNEUROSCI.17-03-00932.1997

57 Schlag, B. D. et al. Regulation of the glial Na+-dependent glutamate transporters by cyclic AMP analogs and neurons. Mol Pharmacol 53, 355–369 (1998). https://doi.org/S0026-895X(24)13190-2 [pii]10.1124/mol.53.3.355

58 Chan, T. J., Her, L. S., Liaw, H. J., Chen, M. C. & Tzeng, S. F. Retinoic acid mediates the expression of glutamate transporter-1 in rat astrocytes through genomic RXR action and non-genomic protein kinase C signaling pathway. J Neurochem 121, 537– 550 (2012). 10.1111/j.1471-4159.2012.07715.x

59 Stenovec, M., Kreft, M., Grilc, S., Pangrsic, T. & Zorec, R. EAAT2 density at the astrocyte plasma membrane and Ca(2 + )-regulated exocytosis. Mol Membr Biol 25, 203–215 (2008). https://doi.org/791039638 [pii]10.1080/09687680701790925

60 Potokar, M. et al. Cytoskeleton and vesicle mobility in astrocytes. Traffic 8, 12–20 (2007). https://doi.org/TRA509 [pii]10.1111/j.1600-0854.2006.00509.x

61 Hughes, E. G., Maguire, J. L., McMinn, M. T., Scholz, R. E. & Sutherland, M. L. Loss of glial fibrillary acidic protein results in decreased glutamate transport and inhibition of PKA-induced EAAT2 cell surface trafficking. Brain Res Mol Brain Res 124, 114– 123 (2004). https://doi.org/S0169328X0400083X [pii]10.1016/j.molbrainres.2004.02.021

62 Choi, H. B. et al. Metabolic communication between astrocytes and neurons via bicarbonate-responsive soluble adenylyl cyclase. Neuron 75, 1094–1104 (2012). https://doi.org/S0896-6273(12)00769-6 [pii]10.1016/j.neuron.2012.08.032

63 Cullum, S. A., Veprintsev, D. B. & Hill, S. J. Kinetic analysis of endogenous beta(2) - adrenoceptor-mediated cAMP GloSensor responses in HEK293 cells. Br J Pharmacol 180, 1304–1315 (2023). https://doi.org/BPH16008 [pii]10.1111/bph.16008

64 Sobolczyk, M. & Boczek, T. Astrocytic Calcium and cAMP in Neurodegenerative Diseases. Front Cell Neurosci 16, 889939 (2022). 10.3389/fncel.2022.889939

