## Supplementary information for "β-Adrenergic regulation of mGluR5-dependent Ca²⁺ signalling in reactive astrocytes"

**Experimental design.** Primary rat cortical astrocytes were cultured in DMEM + 10% FBS (DMEM+) or neurobasal-A + B27 + 5 ng/ml hbEGF (NB+). Prior $t_0$, cells were exposed to extracellular solution containing only 2 mM L-Gln. From $t_0$ to $t_{30\ min}$, cells were exposed to 100 µM L-Glu in addition to 2 mM L-Gln. Measurements for $t_0$ and $t_{30\ min}$ were obtained from separate plates and values normalised to an equivalent final number of 10 000 cells. Each condition comprised four normalised well-level measurements (two wells from 10 000 and two from 5 000 seeded cells).

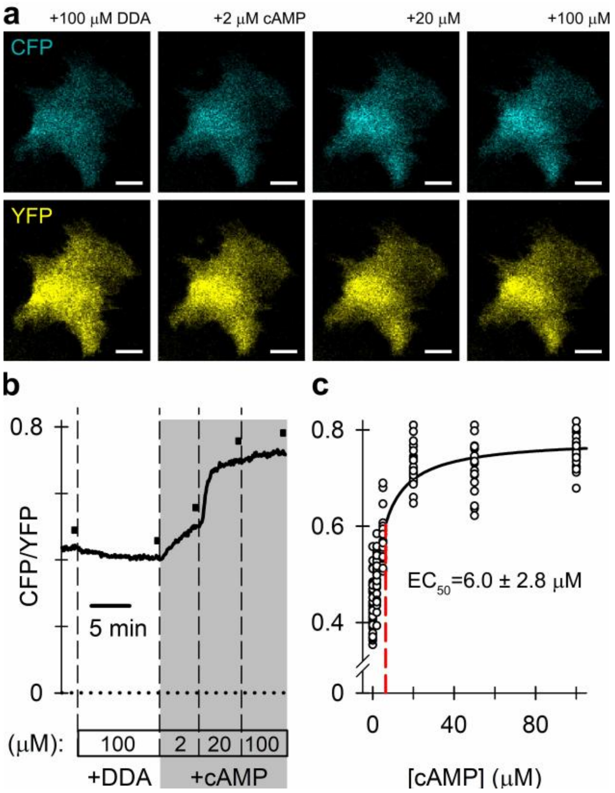


**SUPPLEMENTARY FIGURE 1** Principle of FRET-based fluorescent nanosensor ICUE3 calibration for measurements of astrocyte intracellular cAMP concentration ($[cAMP]_i$). (**a**) Confocal micrographs of transfected DMEM+ astrocytes expressing the fluorescent cAMP nanosensor ICUE3. Emission fluorescence of CFP (top) and YFP (bottom) in ICUE3-transfected DMEM+ astrocytes treated with 100 µM dideoxyadenosine (DDA; an adenylyl cyclase inhibitor), and then with 2, 20 and 100 µM of membrane-permeant, metabolically activable cAMP analogue 8-Br-2'-O-Me-cAMP-AM ("cAMP" above images for simplicity) to increase $[cAMP]_i$. A stepwise increase in $[cAMP]_i$ increased CFP emission while decreasing YFP emission. Scale bars: 20 µm. (**b**) Time-resolved changes in CFP/YFP emission ratio plotted as a function of $[cAMP]_i$. The sequential addition of 100 µM DDA (10 min), followed by 2, 20, and 100 µM membrane-permeant cAMP analogue changed the CFP/YFP emission ratio from 0.44 (resting value) to 0.43, 0.50, 0.58, 0.73, 0.73 and 0.76; each value determined as an average within 30 s (small black rectangles) at the end of the given period (delimited by dashed vertical lines). The grey rectangular area indicates the time of addition of a membrane-permeant cAMP analogue to astrocytes. (**c**) Calibration curve displaying the relationship between cAMP concentration and the CFP/YFP emission ratio in nanosensor ICUE3. Data were fitted by the four parameter logistic curve $f = min + (max-min)/(1 + (x/EC_{50})^{(-Hillslope)2})$, with min = 0.42 ± 0.03, max=0.78 ± 0.05, $EC_{50}$ = 6.02 ± 2.79, and Hill slope = 1.00 ± 0.40.

**SUPPLEMENTARY TABLE 1. Cell-number-normalised metabolites and apparent net accumulation rates**

| Parameter | DMEM+ L-Glu | DMEM+ L-Gln | NB+ L-Glu | NB+ L-Gln |
|---|---|---|---|---|
| **$t_0$ (μM)** | 1.941 ± 0.287 | 3.623 ± 0.425 | 0.344 ± 0.048 | 1.483 ± 0.244 |
| **$t_{30\ min}$ (μM)** | 4.578 ± 0.791 | 4.167 ± 0.612 | 1.549 ± 0.382 | 3.396 ± 0.704 |
| **$t_0$ (pmol/10k cells)** | 116.5 | 217.4 | 20.6 | 89.0 |
| **$t_{30\ min}$ (pmol/10k cells)** | 274.7 | 250.0 | 92.9 | 203.7 |
| **$\Delta t_{30\ min} - t_0$ (pmol/10k cells)** | 158.2 | 32.6 | 72.3 | 114.8 |
| **Apparent net rate (pmol/min/10k cells)** | 5.27 | 1.09 | 2.41 | 3.83 |
| **Fold $t_{30\ min}/t_0$** | 2.36 | 1.15 | 4.51 | 2.29 |
| **P ($t_0$ vs $t_{30\ min}$)** | 0.020 | 0.493 | 0.020 | 0.042 |

**Calculation note.** We obtained concentrations directly from the assay calibration curve. For a 60 µl cell lysate, 1 µM corresponds to 60 pmol per normalised 10 000-cell sample. Apparent net accumulation rates were calculated as $(t_{30\ min} - t_0)/30$ min. These rates describe changes in the intracellular metabolite pool and should not be interpreted as direct transporter flux or direct glutamate-to-glutamine conversion rates.

**SUPPLEMENTARY TABLE 2. Pairwise statistical comparisons.**

| Analyte | Comparison | Test | P value | Significance |
|---|---|---|---|---|
| L-Glu | DMEM+: $t_0$ vs $t_{30\ min}$ | Student unpaired t-test | 0.020 | * |
| L-Glu | NB+: $t_0$ vs $t_{30\ min}$ | Student unpaired t-test | 0.020 | * |
| L-Glu | DMEM+ vs NB+ at $t_0$ | Welch unpaired t-test | 0.010 | * |
| L-Glu | DMEM+ vs NB+ at $t_{30\ min}$ | Student unpaired t-test | 0.014 | * |
| L-Gln | DMEM+: $t_0$ vs $t_{30\ min}$ | Student unpaired t-test | 0.493 | ns |
| L-Gln | NB+: $t_0$ vs $t_{30\ min}$ | Student unpaired t-test | 0.043 | * |
| L-Gln | DMEM+ vs NB+ at $t_0$ | Student unpaired t-test | 0.005 | ** |
| L-Gln | DMEM+ vs NB+ at $t_{30}$ min | Student unpaired t-test | 0.440 | ns |

**Statistical note.** Data for $t_0$ and $t_{30\ min}$ were obtained in separate plates and analysed as unpaired observations. We assessed equal variance for each comparison; we used Welch correction when the equal-variance assumption was not met. Statistical inference is based on normalised well-level measurements from this experiment (n = 4 wells per condition).
